# Wnt signaling rescues amyloid beta induced stem cell loss

**DOI:** 10.1101/2021.09.06.459094

**Authors:** Prameet Kaur, Ellora Hui Zhen Chua, Wen Kin Lim, Nathan Harmston, Nicholas S. Tolwinski

## Abstract

Previously, we established an optogenetic model to induce Amyloid-β intracellular oligomerization to model distinct disease etiologies (Lim *et al*. 2020). Here we examine the effect of Wnt signaling on Amyloid in this model. We observe that Wnt activation rescues the detrimental effects of Amyloid expression and oligomerization. We analyze the gene expression changes downstream of Wnt that contribute to this rescue and find changes in aging related genes, protein misfolding, metabolism and inflammation. We propose that Wnt expression reduces inflammation through repression of Toll activating factors and confirm that chronic Toll activation reduces lifespan. We propose that the protective effect observed for Lithium treatment functions at least in part through Wnt activation and inhibition of inflammation.

## Introduction

Alzheimer’s disease (AD) is an age-related disease affecting millions of people worldwide (D-Paula *et al*. 2012; Kumar *et al*. 2015). No effective therapy exists despite some recent clinical trials and one controversial new drug which has been recently approved (Sevigny *et al*. 2016; Anderson *et al*. 2017; Lalli *et al*. 2021). AD drug development has primarily been based on the amyloid cascade hypothesis, with many drugs targeting amyloid beta (Aβ) directly (Cummings 2018; Cummings *et al*. 2018). The hypothesis postulates that extracellular deposition of Aβ leads to neuronal cell death driving AD (Hardy and Higgins 1992), but whether these plaques are causative remains controversial due to observations of plaques in asymptomatic individuals (Driscoll and Troncoso 2011).

Aβ extracellular aggregates form macroscopic plaques that correlate with cell loss, but Aβ can also form smaller oligomers with soluble oligomers showing the highest toxicity (Ferreira and Klein 2011). We recently described an optogenetic method to study Aβ protein oligomerization *in vivo* in several model organisms (Lim *et al*. 2020). Optogenetics refers to genes that are modified with a light responsive protein domain, allowing spatial and temporal regulation of proteins (Lim *et al*. 2021). Temporal and spatial control is achieved by light exposure of a specific wavelength, without the need to introduce other external agents (Möglich and Moffat 2010; Fenno *et al*. 2011). We used a modified version of the *Arabidopsis thaliana* cryptochrome 2 (CRY2) protein, which clusters in response to blue light (Mas *et al*. 2000).

Our previous studies focused on neurogenesis and metabolism, as well as interventions that could ameliorate the condition (Teo *et al*. 2019; Lim *et al*. 2020). We showed that Lithium worked well to extend the lifespan of *C. elegans* and *Drosophila* in optogenetic Aβ models. We posited that the Wnt signaling pathway may be involved but did not delineate the pathway through which Lithium worked. Here, we present evidence that Wnt signaling can ameliorate the detrimental effects of Aβ oligomerization by promoting stem cell homeostasis and preventing inflammation. We speculate that Wnt activators could function in various tissues affected by amyloid or amyloidosis.

## Results

To study the role of intracellular Aβ we developed a system of expression coupled with light inducible oligomerization (Figure 1a). Previously, we showed a distinction between simple overexpression of Aβ and light induced aggregation where one led to metabolic changes (Teo *et al*. 2019) and the second led to physical damage of the nervous system (Lim *et al*. 2020). In *Drosophila*, transgenic animals expressed the 42-amino-acid human Aβ peptide fused to Cryptochrome 2 and the fluorescent protein mCherry (Aβ^1-42^-CRY2-mCh, Figure 1a). Expression in *Drosophila* used the GAL4/UAS system (Brand and Perrimon 1993) with Elav-Gal4 driving expression in neurons (Video 1 and 2, Figure 1). Glial cells were imaged using the QUAS system with repo-QF2 driving QUAS-GFP (Potter *et al*. 2010). We imaged embryonic neurogenesis in detail and observed physical breakdown of the central nervous system upon Aβ oligomerization (Video 1 and 2, Figure 1b and 1c). In lifespan analysis, we observed a rescue of Aβ-induced lifespan shortening through treatment with Lithium but were unable to introduce Lithium into embryos to test its effect on neurogenesis (Lim *et al*. 2020). We proposed that the rescue may be due to Wnt signaling and attempted to test this in our embryonic system by expressing Wingless (Wg or Wnt1) in developing embryos. However, expression of Wg in the developing nervous system was not successful as embryos ceased to develop (Video 3 and 4, Figure 1d and 1e). In order to establish the effect of Wnt signaling on Aβ activity, we moved to an adult system where Wnt is involved in tissue homeostasis through its ability to maintain stem cells, the *Drosophila* gut.

**Figure 1.**
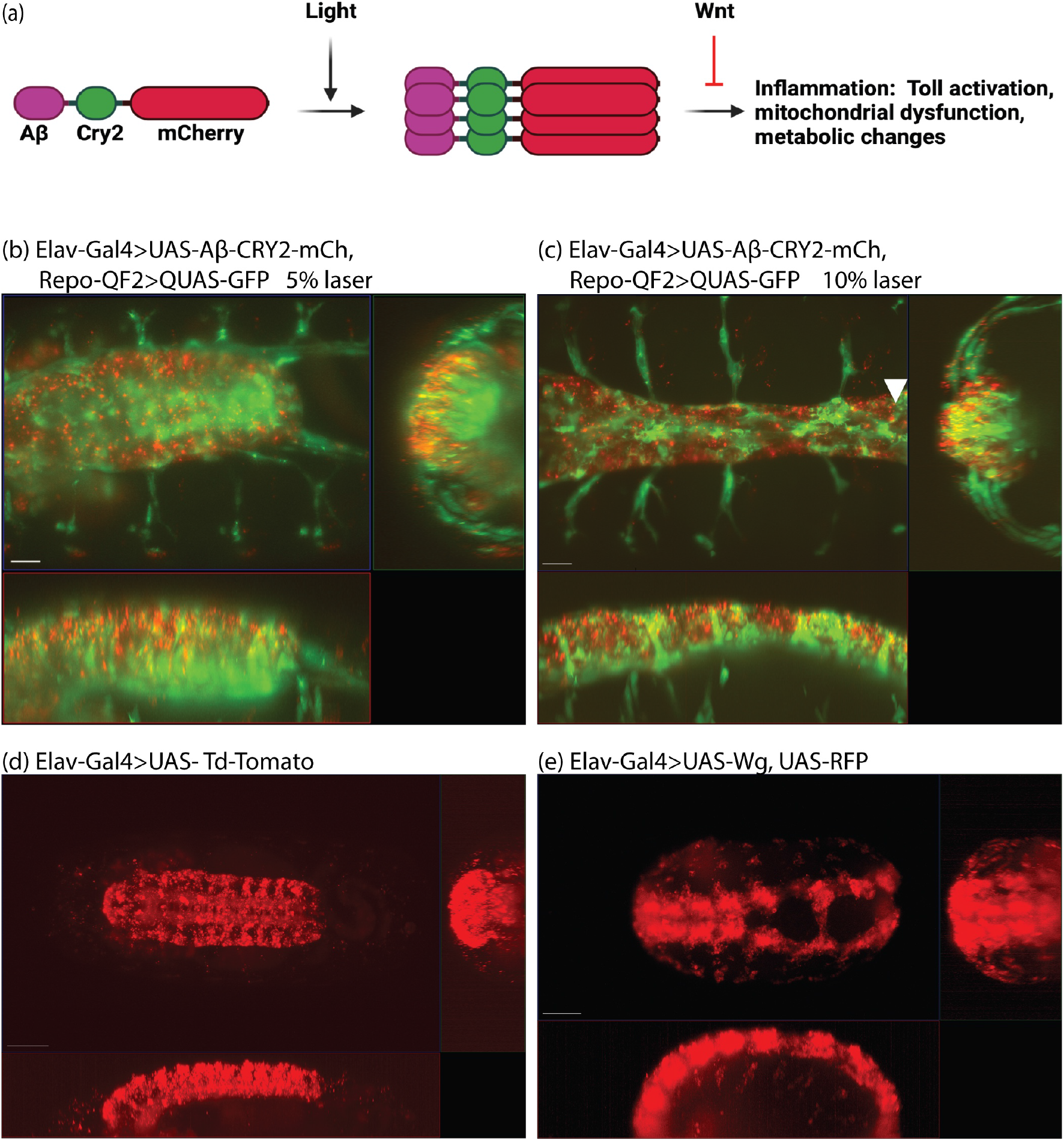
The optogenetic Amyloid system in embryos. (a) Schematic of light-induced Aβ-Cry2mCh clustering in *Drosophila melanogaster* resulting in activation of inflammatory processes which can be rescued by Wnt. Stills from Video 1 (b), Video 2 (c), Video 3 (d), Video 4 (e).

The *Drosophila* gut has rapidly been established as a simple, accessible model for homeostasis, regeneration, and proliferation (Jasper 2020). The tissue is composed of four cell types: enterocytes (ECs or absorptive cells), enteroendocrine (EEs or secretory cells), enteroblasts (EBs or transit amplifying cells) and intestinal stem cells (ISCs). ISCs rest on the external surface of the gut epithelium away from the gut lumen and divide symmetrically to generate more ISCs or asymmetrically to form EBs (Figure 2a) (Zhang and Edgar 2021). We confirmed the presence of gut Wg expression by imaging the midgut where a wg-Gal4 enhancer drove expression of GFP (Figure 2b). As previously shown, expression was most prevalent in the compartment boundaries and stem cell niches (Tian *et al*. 2016; Tian *et al*. 2019). In normal guts, ISCs are sparse and are specifically marked by expression of *escargot*. Therefore *esg* can be used to drive exogenous GAL4 and activate UAS-GFP or another UAS driven gene specifically in ISCs (Micchelli and Perrimon 2006). We expressed Aβ^1-42^-CRY2-mCh in this tissue and imaged the effect on ISCs. In flies kept in the dark where optogenetic oligomerization was inactive, there was a dramatic increase in GFP-positive cells especially in transit amplifying, EBs (Figure 3a-a’’, wildtype guts show an average of five ISCs and no EBs per imaged segment, whereas Aβ^1-42^-CRY2-mCh (Figure 3b-b’’) expressing guts show ~9 ISCs and ~24 EBs per imaged segment). This effect was completely abrogated by the co-expression of Wg along with Aβ^1-42^-CRY2-mCh where we observed an increase in total ISCs, but no increase in EBs (Figure 3c-c’’, Aβ^1-42^-CRY2-mCh + Wg expressing guts showed ~14 ISCs and ~2 EBs). We compared the effect on flies kept in the dark (no optogenetic oligomerization) to flies exposed to light (blue light induced CRY2-based oligomerization). We observed a small increase in GFP-positive EBs (Figure 3d-d’’, Aβ^1-42^-CRY2-mCh + Light expressing guts showed on average 5 ISCs and 34 EBs) and a clear loss of EBs and an increase in the number of ISCs following co-expression of Wg along with Aβ^1-42^-CRY2-mCh (Figure 3e-e’’, Aβ^1-42^-CRY2-mCh + Wg + Light expressing guts showed on average 22 ISCs and 2 EBs). Expression of Wg alone led to an increase in ISC numbers (Figure 3f-f’’, Wg expressing guts showed on average 15 ISCs and 2 EBs). Together, these findings suggest that either overexpression or light-induced oligomerization of Aβ^1-42^-CRY2-mCh led to asymmetric stem cell divisions and dramatically altered tissue homeostasis. But most importantly, tissue homeostasis was restored following overexpression of Wg.

**Figure 2.**
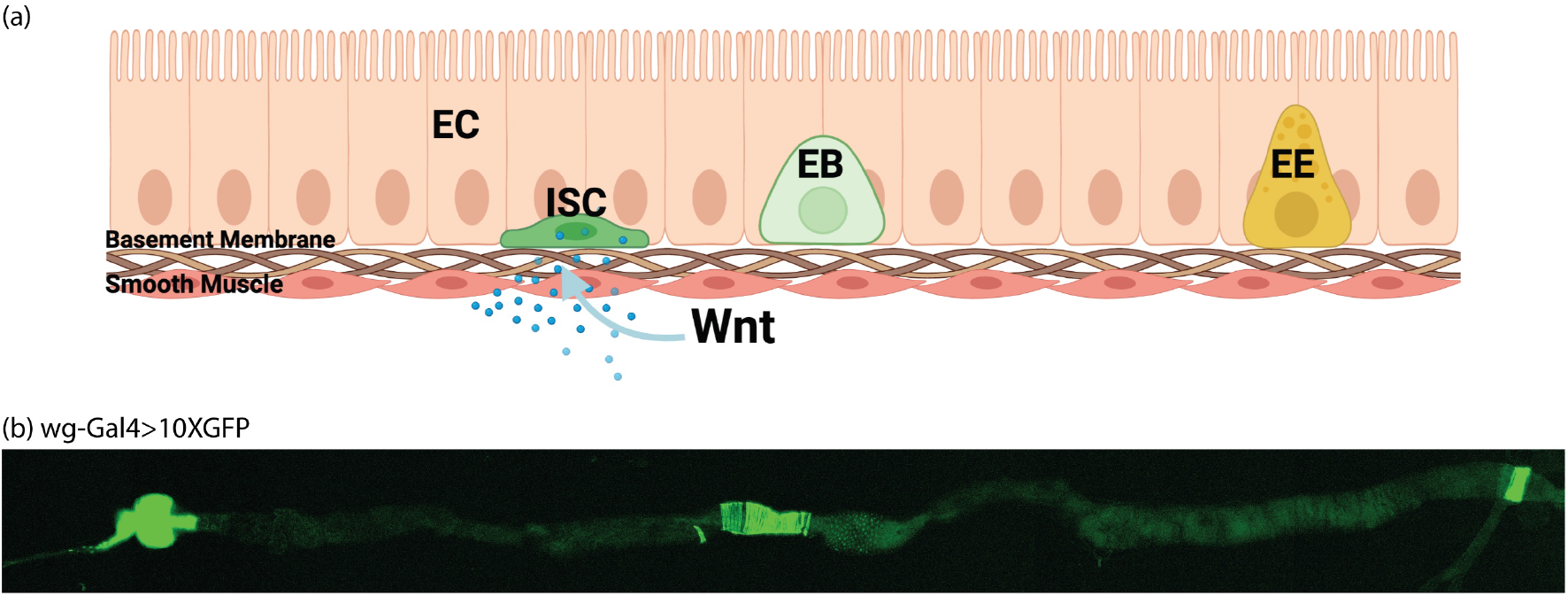
Wnt expression in the *Drosophila* gut. (a) Model of the *Drosophila* gut comprising enterocytes (ECs), enteroendocrine (EEs), enteroblasts (EBs) and intestinal stem cells (ISCs). ISCs rest on the external surface of the gut epithelium away from the gut lumen and divide symmetrically to make more ISCs or asymmetrically to form EBs. (b) representative midgut section of a wg-Gal4>10XGFP fly.

**Figure 3.**
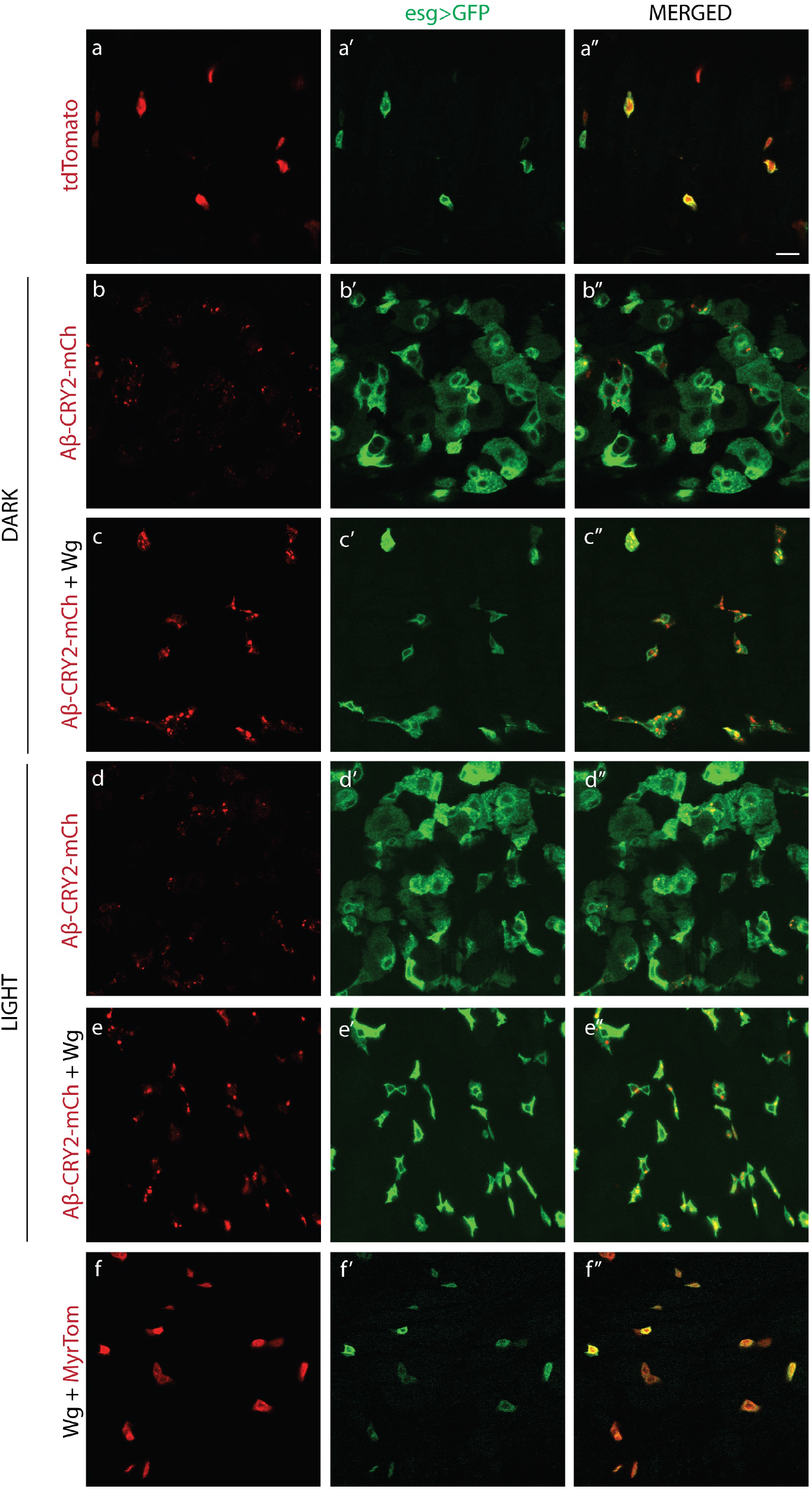
Wnt and Amyloid expression in ISCs. Images of midgut sections of (a-a’’) esg-Gal4>UAS-GFP, UAS-Td-Tomato flies, (b-b’’) esg-Gal4>UAS-GFP, UAS-Aβ^1-42^-CRY2-mCh flies kept in the dark, (c-c’’) esgGal4>UAS-GFP, UAS-wg, UAS-Aβ^1-42^-CRY2-mCh flies kept in the dark, (d-d’’) esg-Gal4>UAS-GFP, UAS-Aβ^1-42^-CRY2-mCh flies kept in the light, (e-e’’) esg-Gal4>UAS-GFP, UAS-wg, UAS-Aβ^1-42^-CRY2-mCh flies kept in the light and (f-f’’) esg-Gal4>UAS-GFP, UAS-wg, UAS-myr-Tomato flies. Scale bar represents 10μm.

Expression of Aβ^1-42^-CRY2-mCh in whole organisms or specifically in the nervous system led to decreased life and health spans in a light dependent manner (Lim *et al*. 2020). We postulated that the loss of homeostasis observed in the *Drosophila* gut would be detrimental to adult survival, and so tested the lifespans of flies expressing amyloid and Wg in ISCs. We observed a significant decrease in survival of flies expressing Aβ^1-42^-CRY2-mCh grown in the light (Figure 4). This effect was rescued by expression of Wg alongside. Wg expression alone was detrimental as well, but the effect was equivalent to Aβ^1-42^-CRY2-mCh combined with Wg suggesting that Wg rescued the Aβ^1-42^-CRY2-mCh effects, but still led to other detriments.

**Figure 4.**
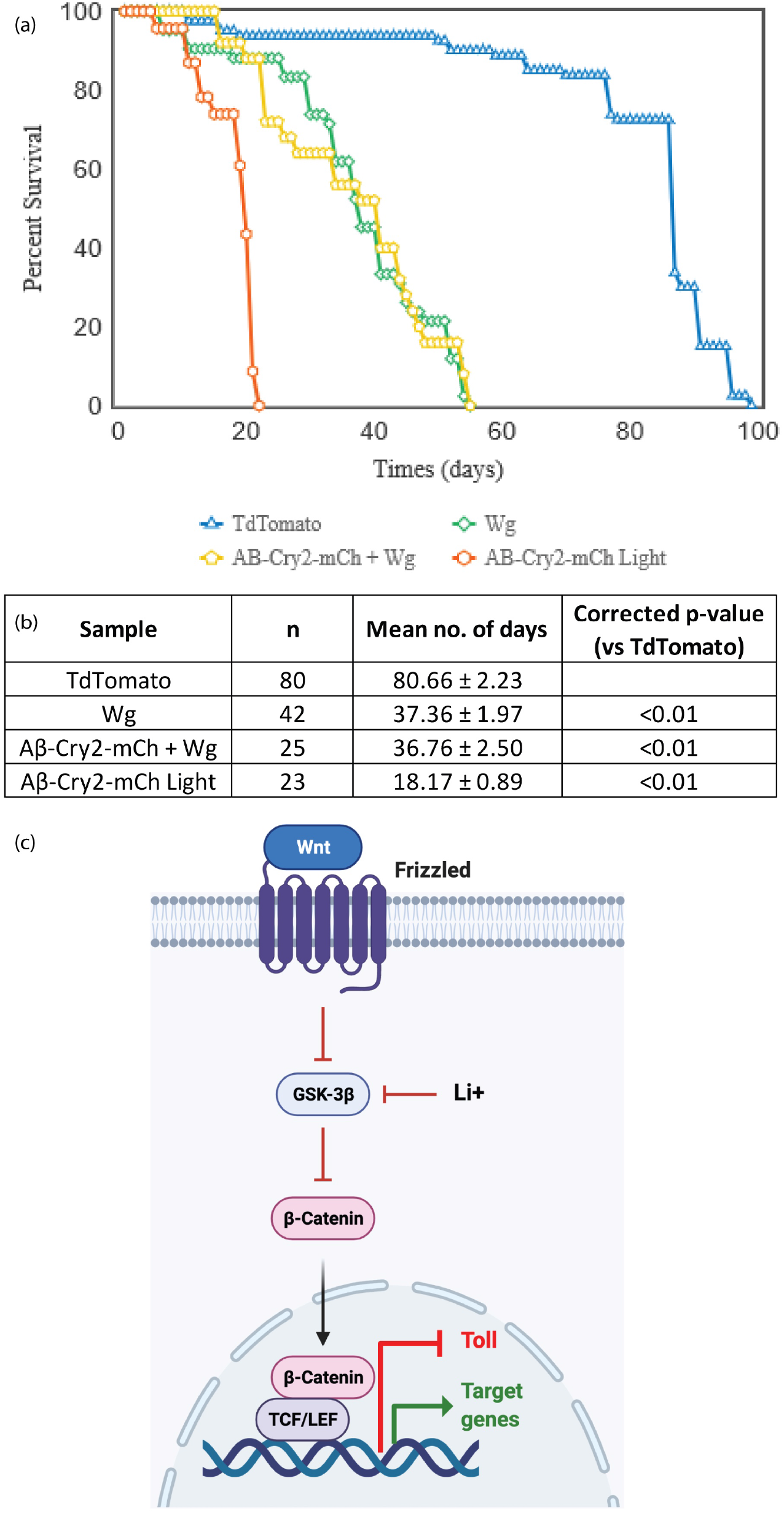
Lifespan decrease rescued by Wnt expression. Lifespan analysis of flies expressing TdTomato (control), Wg, Aβ-CRY2-mCh + Wg and Aβ-CRY2-mCh in gut stem cells. Genotype: TdTomato – esg-Gal4>UAS-GFP, UAS-Td-Tomato; Wg – esg-Gal4> UAS-GFP, UAS-wg; Aβ-CRY2-mCh + Wg – esg-Gal4> UAS-GFP, UAS-wg, UAS- Aβ^1-42^-CRY2-mCh; Aβ-CRY2-mCh – esg-Gal4> UAS-GFP, UAS- Aβ^1-42^-CRY2-mCh. (c) Schematic of Lithium induced activation of the Wnt pathway to inhibit Toll signalling.

The finding that Wg rescued both the homeostatic and lifespan deficits caused by Aβ expression in ISCs strongly supports our hypothesis that the inhibition of Glycogen Synthase Kinase 3 (GSK-3) by Lithium, leading to the subsequent activation of the Wnt pathway, was responsible for the observed differences in survival (Figure 4c) (Lim *et al*. 2020). To identify the potential pathways and mechanisms responsible for this rescue, we collected *Drosophila* guts from flies exposed to light, expressing Aβ^1-42^-CRY2-mCh (Aβ), Wg alone (Wg), Aβ^1-42^-CRY2-mCh and Wg together (WgAβ), or control flies (WT) expressing only a fluorescent protein, and performed RNA-seq (Figure 5a). Our analysis identified 2800 genes (FDR < 10%) as significantly differentially expressed across the conditions (Figure 5b, Supplemental Table 1). Clustering of these genes identified six distinct clusters, each representing groups of genes with similar expression profiles across the four conditions investigated (Figure 5b, c).

**Figure 5.**
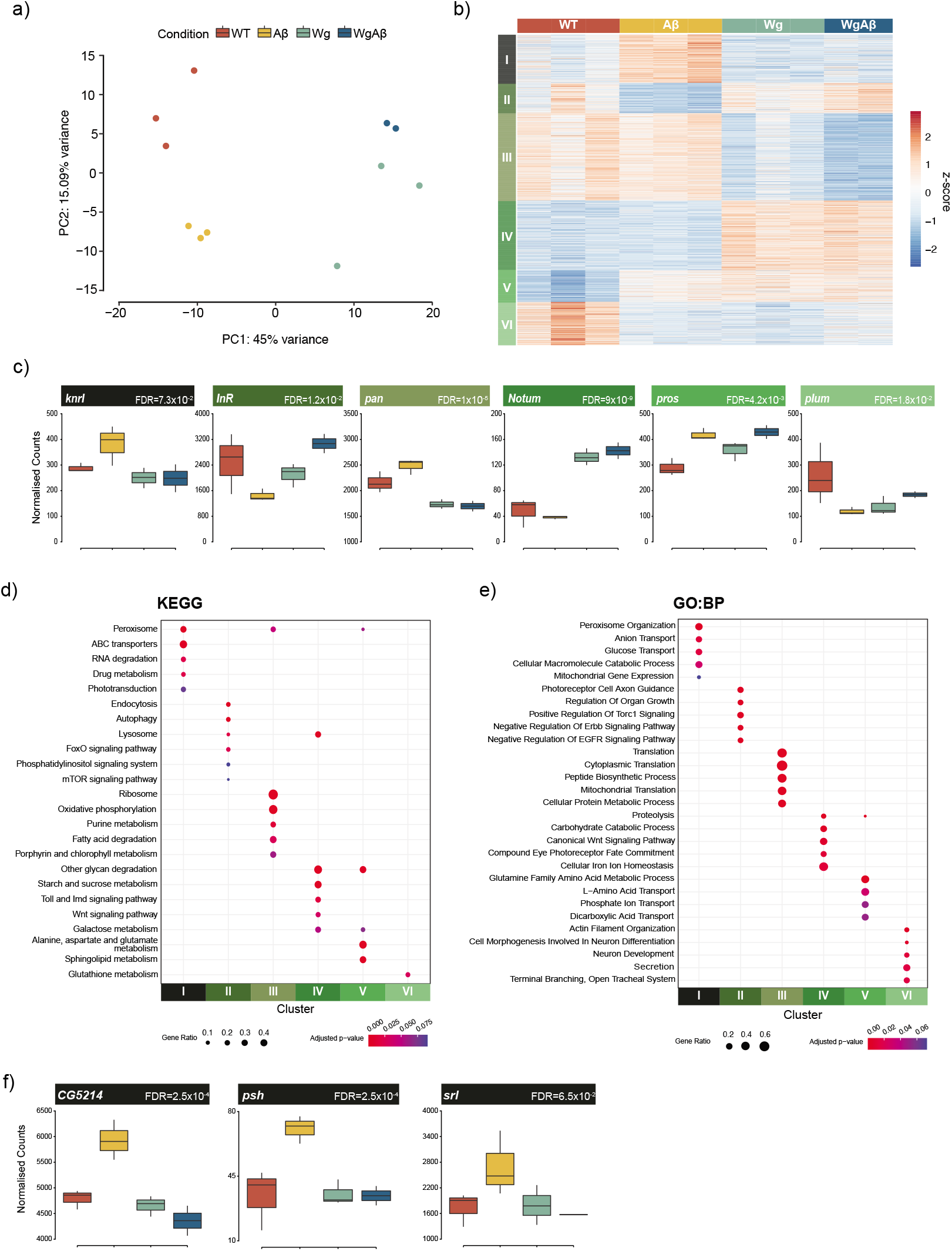
Gene expression changes in response to Amyloid and Wnt. (a) Principal components analysis of RNA-seq data reveals clear separation of samples by experimental condition (b) Clustering of differentially expressed genes identifies six clusters with distinct expression patterns across the four conditions (c) Expression profiles of representative genes for each of the six clusters (d) KEGG pathway enrichments identifies key developmental and homeostatic pathways associated with the individual clusters (FDR < 10%) (e) Biological process (GO:BP) analysis identifies distinct processes enriched in each of the clusters (FDR < 10%). (f) Expression profiles of genes from Cluster I which are relevant to AD biology.

Each of the six clusters was enriched for distinct pathways and processes (Figure 5d, e, Supplemental Tables 2 and 3). Cluster I (N = 440) contained genes that were upregulated by expression of Aβ^1-42^-CRY2-mCh but were repressed by increased Wnt signaling. Genes in this cluster were associated with peroxisome, transport, and metabolic processes (Supplemental Figure S1). Multiple studies have previously directly implicated the peroxisome in mediating AD pathology (Kou *et al*. 2011), and have implicated metabolic changes in the disease (Jo *et al*. 2020). Conversely, the genes in Cluster II (N = 273) consisted of genes repressed by Aβ^1-42^-CRY2-mCh, but whose expression was rescued following activation of Wnt signaling. This cluster was enriched for processes associated with autophagy, lysosomal activity (Supplemental Figure S2), and the regulation of signaling pathways whose activation has previously been found to be protective against AD, e.g. EGFR, FoxO, and mTOR (Oddo 2012; Cai *et al*. 2015; Suresh *et al*. 2020). Cluster III (N = 785) and Cluster IV (N = 630) consisted of genes that were Wnt-repressed or Wnt-activated respectively, *i.e*. were up- or down-regulated following overexpression of Wg, but expression of Aβ^1-42^-CRY2-mCh had no effect on their expression. Cluster III was enriched for ribosome and translation-related processes, whereas Cluster IV was enriched for the Wnt signaling pathway and contained known Wnt target genes (*e.g. Notum)*. Cluster V (N = 285) represented genes that were downregulated in the WT condition but were upregulated in all other conditions, while the genes in Cluster VI (N = 387) displayed the opposite pattern. It is likely that these two clusters simply represent the effects of perturbing the system. Clusters I and II represent groups of genes whose expression is affected by expression of Aβ and is either repressed or rescued by activation of Wnt signaling and likely contain genes responsible for the observed differences in lifespan.

Within Cluster I, we identified several potential targets (Figure 5f), including CG5214, a succinyltransferase that regulates post-translational modifications of proteins and is associated with aging (Hong *et al*. 2016; Bonnay *et al*. 2020), and the transcriptional coactivator spargel (*srl*), the *Drosophila* homolog of *PGC-1α*, a gene involved in mitochondrial homeostasis and Insulin-TOR signaling, which has been implicated in AD pathogenesis (Tiefenböck *et al*. 2010; Mukherjee and Duttaroy 2013). We further observed genes related to innate immune activation, such as Persephone (*psh*), a serine protease that is involved in regulating Toll pathway activation (Ligoxygakis *et al*. 2002). Overall, our transcriptomic analysis showed Wnt rescuing the expression of components of key neuroprotective pathways, while acting against most of the known processes involved in AD pathogenesis.

In AD, inflammation plays an important role in disease progression (Kinney *et al*. 2018). The upregulation of *psh* in Aβ is indicative of activation of the Toll pathway, a key pathway involved in inflammation and innate immunity. We used the Toll receptor, a key activator of innate immune response, to assay the inflammatory activity as it relates to lifespan. We used an optogenetic version of Toll we had previously constructed and tested its effect on fly lifespans (Bunnag *et al*. 2020). We expressed Toll-Cry2-mCherry in adult flies using the esg-Gal4 system to target ISCs and in the whole fly using Arm-Gal4. Whole animal expression of Toll-Cry2-mCherry along with light exposure led to rapid death (Figure 6). Expression only in ISCs was less damaging, but activation of the Toll pathway by Toll-Cry2-mCherry led to a decrease in lifespan, comparable to the effect of Aβ^1-42^-CRY2-mCh. The effects were comparable when tested in flies kept in the dark. Overall, modulating the level of innate immune response led to lifespan shortening.

**Figure 6.**
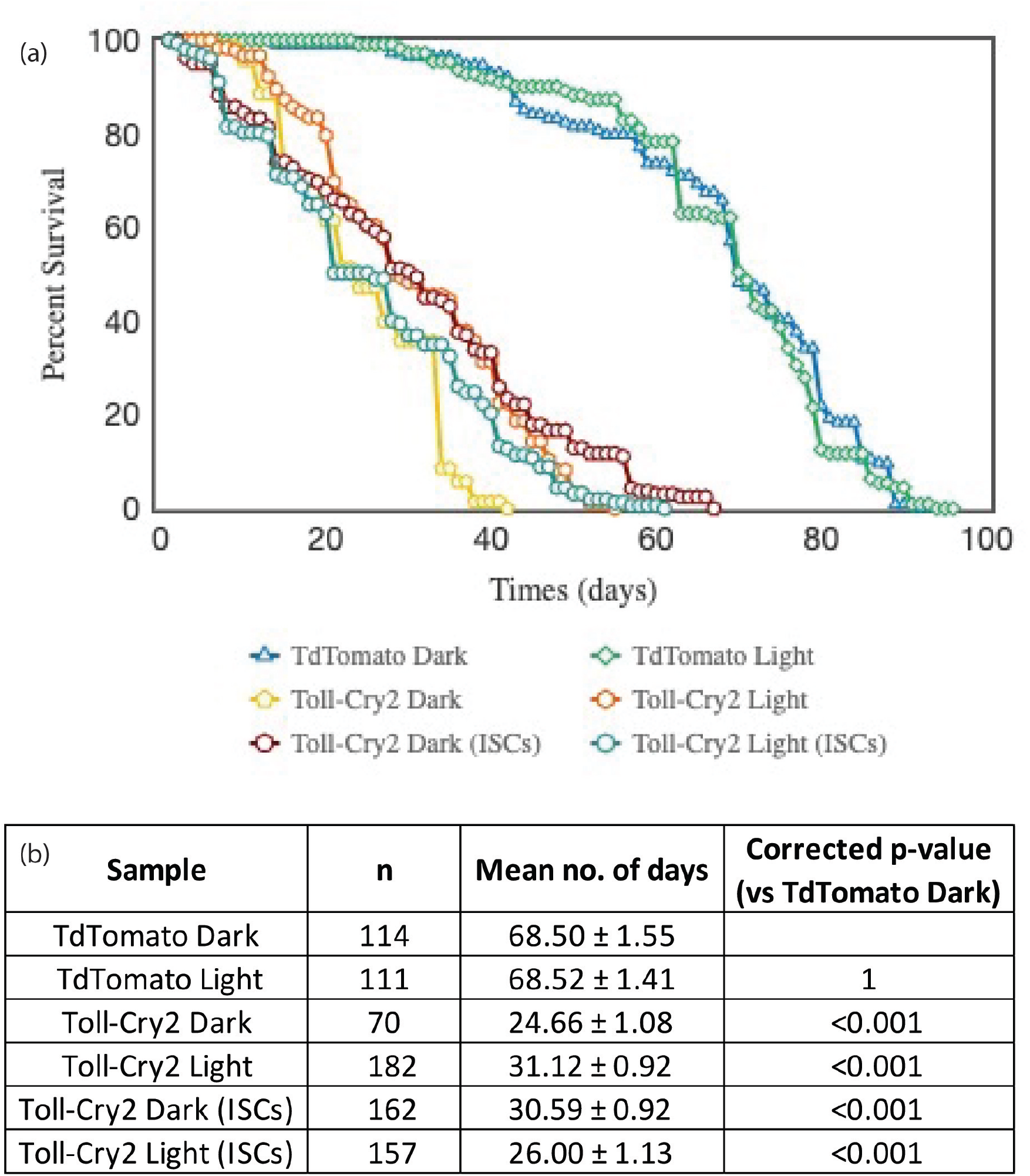
Activated Toll reduces lifespan. Lifespan analysis of TdTomato (control) dark and light, Toll-Cry2-mCh dark and light in the whole fly, Toll-Cry2-mCh dark and light expression in the ISCs only. Genotype: TdTomato – (Arm-Gal4, UAS-GFP), UAS-Td-Tomato; Toll-Cry2 – ArmGal4> UAS-Toll-Cry2-mCh; Toll-Cry2 (ISCs) – esg-Gal4> UAS-GFP, UAS-Toll-Cry2-mCh.

## Discussion

In an update to our previously published study, we followed up on a comment made by one of the referees, namely which mechanism does Lithium use to extend the lifespan of *Drosophila* expressing Aβ^1-42^-CRY2-mCh. We postulated that the Wnt pathway was the key component to this based on previous studies (Ng *et al*. 2019; Suresh *et al*. 2020). However, we were unable to test the model in embryonic neurogenesis as application of Lithium and Wnt to embryos proved either technically or genetically problematic, and instead we investigated this in the *Drosophila* gut, an adult homeostasis model. Overexpression of Wnt in ISCs was found to ameliorate the homeostatic and lethal consequences of Aβ^1-42^-CRY2-mCh expression. We found several metabolic, proteostatic and inflammatory pathways to be activated by Aβ^1-42^-CRY2-mCh and repressed by Wnt. We propose that Wnt’s role in promoting homeostasis in stem cells can be extended to prevent some of the detrimental effects of Aβ^1-42^-CRY2-mCh expression by preventing Toll pathway activation. We show that Toll activation is just as detrimental as Aβ^1-42^-CRY2-mCh expression and propose that Wnt prevents Toll hyperactivation leading to amelioration of amyloid deposition.

As we and others previously proposed that Lithium could be used as a treatment for AD (Sofola-Adesakin *et al*. 2014), by narrowing the effect of Lithium to Wnt activation, this opens the possibility of using Wnt activating drugs in AD treatment, especially as one has recently been shown to be life extending (Castillo-Quan *et al*. 2019). Most importantly, the homeostatic role of Wnt is not the only developmental signal that promotes homeostasis and prevents amyloid detriments. Recently, a role for Hedgehog was discovered in glial cells to promote lifespan and prevent amyloid-dependent neurodegeneration (Rallis *et al*. 2020). These findings together suggest that stem cell homeostasis and re-activation of developmental pathways may be key mechanisms that could be targeted to prevent neurodegeneration.

## Methods

### Crosses and expression of UAS construct

Optogenetic transgenes were generated as previously described in (Kaur *et al*. 2017; Lim *et al*. 2020). Expression was driven by Elav-GAL4 the neuronal driver, *Escargot-GAL4* (Brand and Perrimon 1993) and tubP-GAL80^ts^ (McGuire *et al*. 2003). All additional stocks were obtained from the Bloomington *Drosophila* Stock Center (NIH P40OD018537) that were used in this study.

Fly lines:

Elav-Gal4: BDSC 458 (Lin and Goodman 1994)
Repo-QF2: BDSC 66477 (Potter *et al*. 2010)
QUAS-6XGFP: BDSC 52263 (Shearin *et al*. 2014)
UAS-Aβ-CRY2-mCherry: (Lim *et al*. 2020)
UAS-mCD8.RFP: BDSC 32220 (Pfeiffer *et al*. 2010)
UAS-wg: BDSC 5918 (Hays *et al*. 1997)
UAS-Td-Tomato: BDSC 36328 (Joost Schulte and Katharine Sepp)
Tl(CRISPaint.T2A-GAL4)wg: BDSC #83627 (Bosch *et al*. 2020)
UAS-10XGFP: BDSC 32185 (Pfeiffer *et al*. 2010)
Esg-Gal4, UAS-GFP; tub-Gal80^ts^, UAS-dCas9.VPR: BDSC 67069 (Micchelli and Perrimon 2006)
UAS-myr::tdTomato: BDSC 32222 (Pfeiffer *et al*. 2010)
UAS-Toll-Cry2-mCherry: (Bunnag *et al*. 2020)
Arm-Gal4: BDSC 1560 (White and Vincent 1996)
ArmGal4; tub-GAL80^ts^: BDSC 86327 (McGuire *et al*. 2003)

Fly crosses performed were:

1. Elav-Gal4; Repo-QF2, QUAS-GFP x UAS-Aβ-CRY2-mCh
2. Elav-Gal4, UAS-RFP x UAS-wg
3. Elav-Gal4 x UAS-Td-Tomato
4. Tl(CRISPaint.T2A-GAL4)wg X UAS-10XGFP
5. w; UAS-TdTomato X esg-Gal4, UAS-GFP; tubP-GAL80^ts^
6. w; UAS-Aβ^1-42^-CRY2-mCh X esg-Gal4, UAS-GFP; tubP-GAL80^ts^
7. w; UAS-wg; UAS-Aβ^1-42^-CRY2-mCh X esg-Gal4, UAS-GFP; tub-GAL80^ts^
8. w; UAS-wg, UAS-myr-Tomato X esg-Gal4, UAS-GFP; tub-GAL80^ts^
9. w; UAS-wg X esg-Gal4, UAS-GFP; tub-GAL80^ts^
10. w; UAS-Toll-Cry2-mCh X esg-Gal4, UAS-GFP; tub-GAL80^ts^
11. w; UAS-Toll-Cry2-mCh X ArmGal4; tub-GAL80^ts^
12. w; UAS-Td-Tomato X esg-Gal4, UAS-GFP; tub-GAL80^ts^
13. w; UAS-Td-Tomato X ArmGal4; tub-GAL80^ts^

### Light-sheet microscopy

Embryos at stage 11 were selected using halocarbon oil (Sigma), dechorionated and mounted into the Lightsheet Z.1 (Carl Zeiss, Germany) microscope and imaged with a 40x W Plan-Apochromat 40x.1.0 UV–VIS detection objective (Kaur *et al*. 2018; Kaur *et al*. 2020). Image data were processed using the maximum intensity projection function of ZEN 2014 SP software (Carl Zeiss, Germany).

### Gut preparations and fluorescence microscopy

Adult fly midguts from at least 3 flies were dissected and imaged at the 25^th^ percentile from the anterior midgut on the Zeiss LSM800 (Carl Zeiss, Germany) as described by Bunnag et al., 2020. Images were processed using the ZEN 2014 SP1 software (Carl Zeiss, Germany). Image quantification was done in ImageJ (Schneider *et al*. 2012).

### RNA preparation and RNA-sequencing

Flies were dissected in 1 × PBS. RNA was isolated from at least 10 midguts using the Isolate II RNA Mini kit (Bioline, UK). The extracted RNA was quantified using Nanodrop (Thermo Fisher Scientific). Library preparation was performed using 1μg of total RNA and sequencing was performed using Illumina HiSeq 4000 System (2 × 151 bp read length, 40 million reads per sample) by NovogeneAIT Genomics (Singapore).

### RNA-seq analysis

#### Data processing and QC

RNA-seq was aligned against BDGP6.22 (Ensembl version 97) using STAR v2.7.1a (Dobin *et al*. 2013) and quantified using RSEM v1.3.1 (Li and Dewey 2011). Reads mapping to genes annotated as rRNA, snoRNA, or snRNA were removed. Genes which have less than 10 reads mapping on average across all samples were also removed. Differential expression analysis was performed using DESeq2 (Love *et al*. 2014). The Likelihood Ratio Test (LRT) was used to identify any genes that show change in expression across the different conditions. Pairwise comparisons were performed using a Wald test, with independent filtering. To control for false positives due to multiple comparisons in the genome-wide differential expression analysis, the false discovery rate (FDR) was computed using the Benjamini-Hochberg procedure. The gene-level counts were transformed using a regularized log transformation, converted to z-scores, and clustered using Partitioning Around Medoids (PAM) using correlation distance as the distance metric.

#### Functional Enrichment analysis

Gene Ontology (GO) and KEGG pathway enrichments for each cluster were performed using EnrichR (Chen *et al*. 2013; Kuleshov *et al*. 2016; Xie *et al*. 2021). Terms with an FDR < 10% were defined as significantly enriched.

### Lifespan studies

*Drosophila* were counted daily for the number of dead subjects and the number of censored subjects (excluded from the study). *Drosophila* that failed to respond to taps were scored as dead and those stuck to the food were censored. Lifespan analysis was performed using OASIS 2 (Online Application for Survival Analysis 2) (Han *et al*. 2016). Raw numbers available for figure 4 in Supplementary Table 4 and for figure 6 in Supplementary Table 5.

## Code availability

All code necessary to recreate the results from the RNA-seq analysis is available from: https://github.com/harmstonlab/Ab

## Data availability

RNA-seq data generated from this study has been deposited to GEO (GSE181844)

## Acknowledgements

We are thankful for the funding provided by AcRF grants IG19-SI102 and IG20-BG101 to NST. NH and EHZC are supported by a Yale-NUS and Duke-NUS startup grant.

---

**Video 1.** Elav-Gal4>UAS-Aβ^1-42^-CRY2-mCh, Repo-QF2>QUAS-GFP embryos kept in the dark and imaged for neurons using mCherry and glial cells using low (5%) blue laser power to allow imaging of glial cells without inducing pathological clustering. Scale bar represents 50μm.

**Video 2.** Elav-Gal4>UAS-Aβ^1-42^-CRY2-mCh, Repo-QF2>QUAS-GFP embryos kept in the light and imaged for neurons using mCherry and glial cells using high (10%) blue laser power to allow imaging of glial cells to induce clustering. Scale bar represents 50μm.

**Video 3.** Embryos expressing Td-Tomato in the nervous system (Elav-Gal4>UAS-Td-Tomato) develop normally. Scale bar represents 20μm.

**Video 4.** Expressing wg in the nervous system (Elav-Gal4>UAS-wg, UAS-RFP) is lethal to the embryos. Scale bar represents 20μm.

## Supplemental Figures

**S1:**
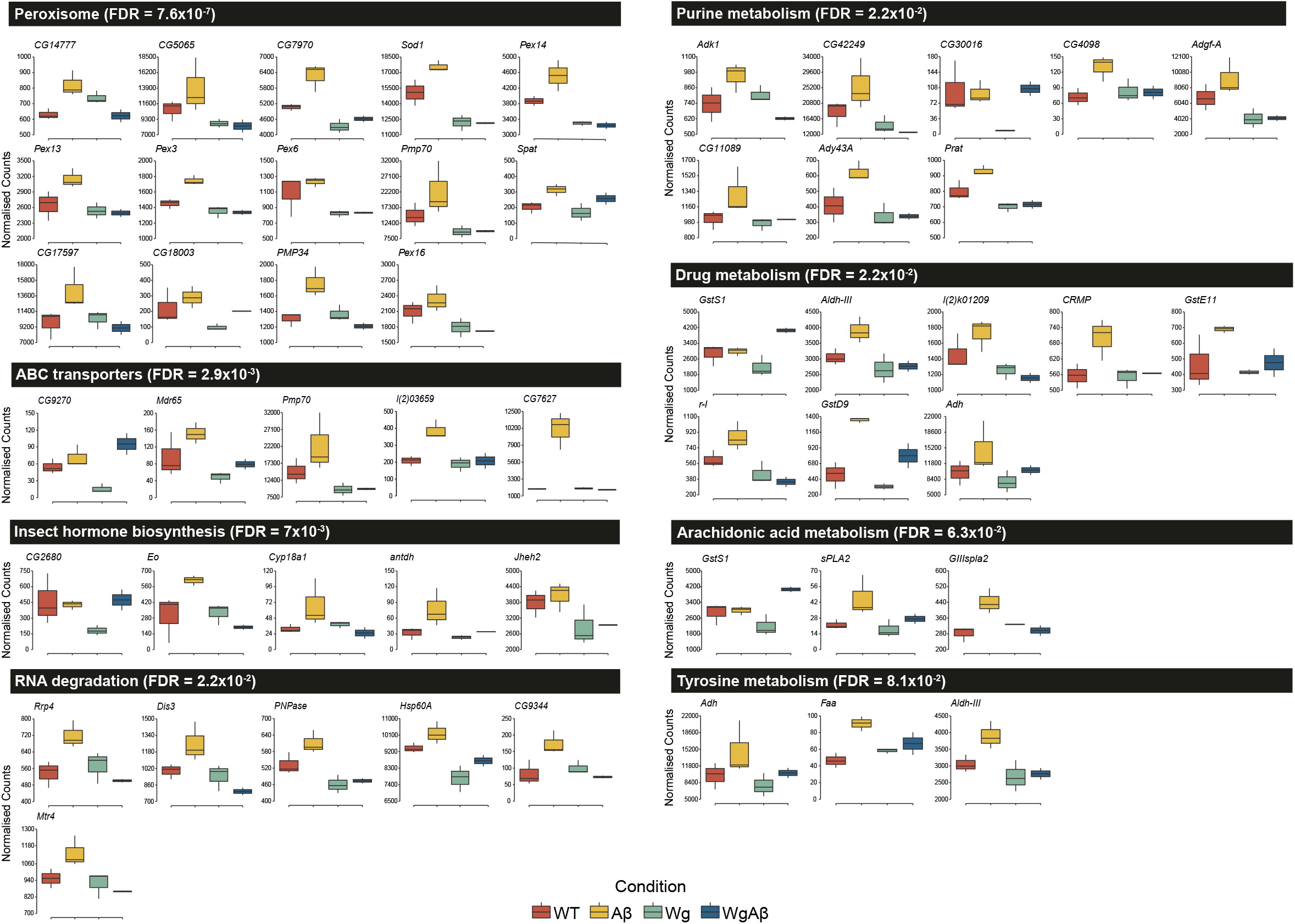
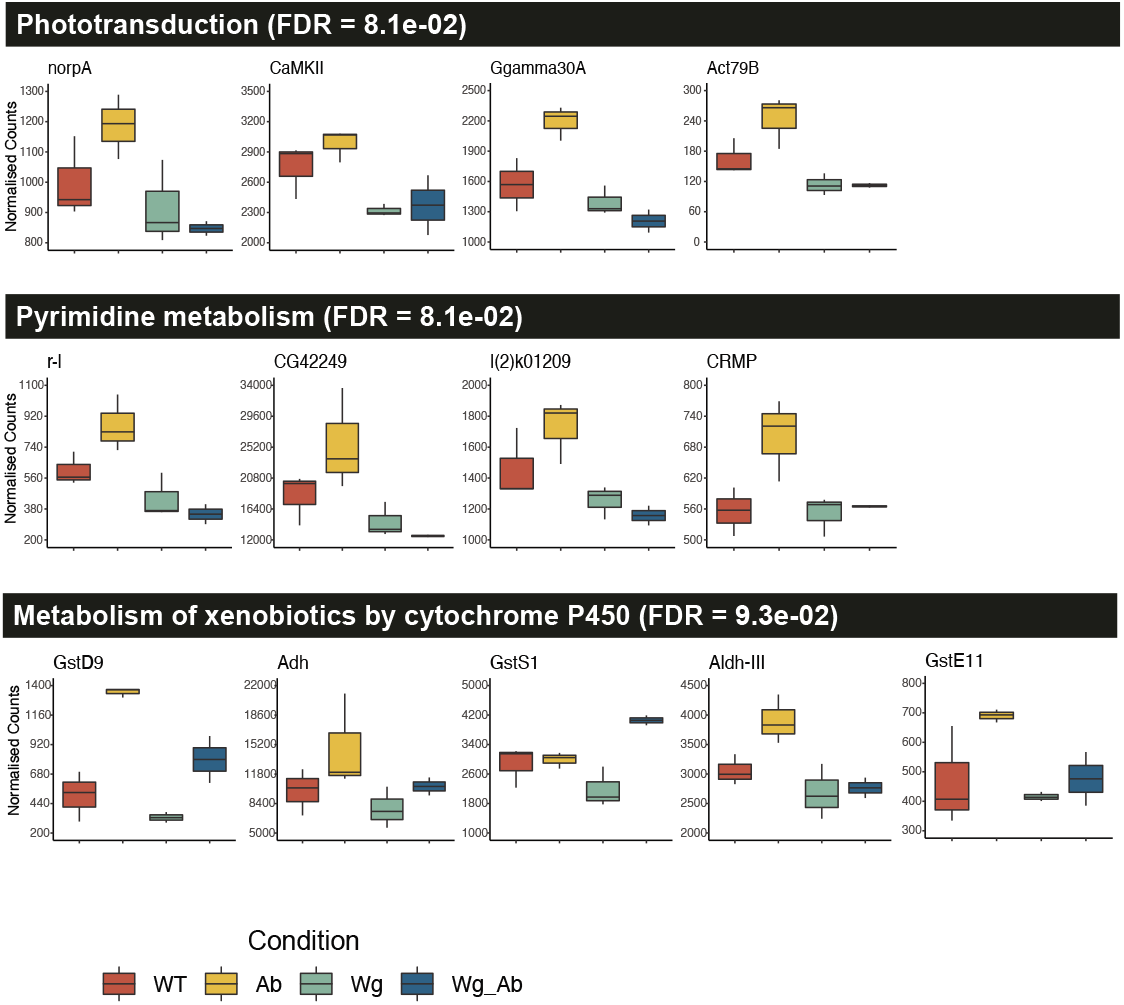
Expression profiles of genes contributing to enrichment of key processes/pathways identified as enriched in Cluster I.

**S2:**
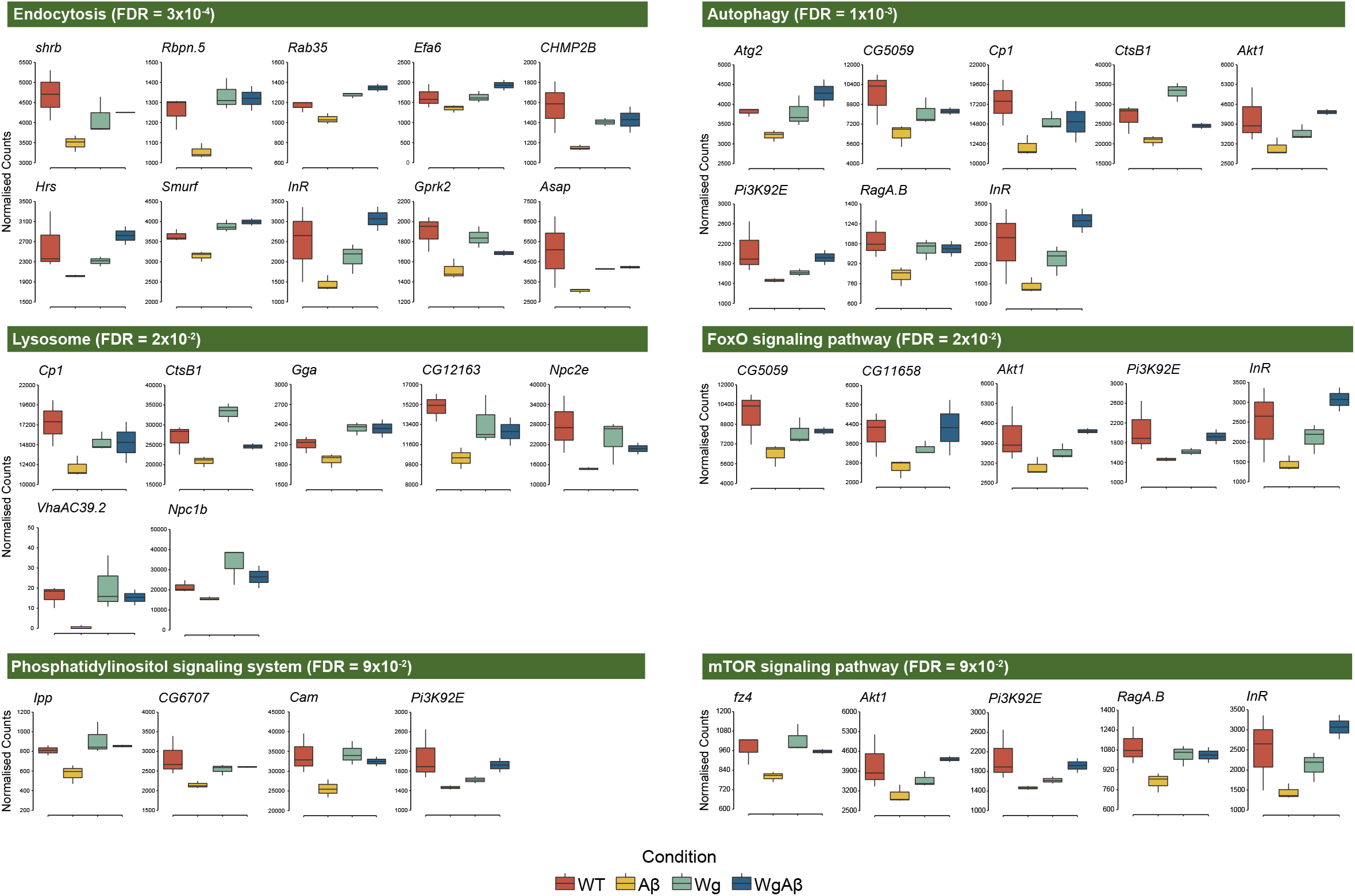
Expression profiles of genes contributing to enrichment of key processes/pathways identified as enriched in Cluster II.

## Supplemental Tables

Table S1: Results from differential expression analysis

Table S2: GO:BP enrichment analysis results for each cluster

Table S3: KEGG pathway enrichment analysis results for each cluster

Table S4: Lifespan Analysis for Wnt and Amyloid

Table S5: Lifespan Analysis for Toll

